# Tumor Characterization using Unsupervised Learning of Mathematical Relations within Breast Cancer Data

**DOI:** 10.1101/2020.06.08.140723

**Authors:** Cristian Axenie, Daria Kurz

## Abstract

Despite the variety of imaging, genetic and histopathological data used to assess tumors, there is still an unmet need for patient-specific tumor growth profile extraction and tumor volume prediction, for use in surgery planning. Models of tumor growth predict tumor size and require tumor biology-dependent parametrization, which hardly generalizes to cope with tumor variability among patients. In addition, the datasets are limited in size, owing to the restricted or single-time measurements. In this work, we address the shortcomings that incomplete biological specifications, the inter-patient variability of tumors, and the limited size of the data bring to mechanistic tumor growth models and introduce a machine learning model capable of characterizing a tumor, namely its growth pattern, phenotypical transitions, and volume. The model learns without supervision, from different types of breast cancer data the underlying mathematical relations describing tumor growth curves more accurate than three state-of-the-art models on three publicly available clinical breast cancer datasets, being versatile among breast cancer types. Moreover, the model can also, without modification, learn the mathematical relations among, for instance, histopathological and morphological parameters of the tumor and, combined with the growth curve, capture the (phenotypical) growth transitions of the tumor from a small amount of data. Finally, given the tumor growth curve and its transitions, our model can learn the relation among tumor proliferation-to-apoptosis ratio, tumor radius, and tumor nutrient diffusion length to estimate tumor volume, which can be readily incorporated within current clinical practice, for surgery planning. We demonstrate the broad unsupervised learning and prediction capabilities of our model through a series of experiments on publicly available clinical datasets.

## 1 Background

With 71888 new cases reported in 2018 in Germany, breast cancer represents 25% of all cancer types affecting the population [1]. Breast cancer assessment has transitioned to novel techniques including, mammography, Magnetic Resonance Imaging (MRI), ultrasound, and optical tools, which are becoming increasingly accessible and affordable [18]. Yet, when considering, for instance, Ductal Carcinoma In Situ (DCIS) [2] – a significant precursor to invasive breast cancer – typical mammogram diagnosis is not accurate (i.e. initial cancer cells typically classified as microcalcifications). This is usually caused by the limited understanding of DCIS growth [9,10], its phenotype which is determined by genomic/proteomic- and microenvironment-dependent stochastic processes [16,19], and its cell volume changes during proliferation and necrosis [6]. The current landscape shows that there is still an unmet need for patient-specific tumor characterization and tumor volume prediction, for use in surgery planning [4,7]. As there is a difference between, for instance, mammography and histopathology estimated sizes, the clinician cannot obtain a “fixed surgical size” of a tumor to be excised. This may contribute to over-treatment, including needless surgery. Only limited work has been done towards patient-calibrated modelling and predictions of tumor clinical progression [15] or patient-specific assessment of surgical volume [8]. Our work addresses this need and finds motivation in the following clinically-pertinent scientific questions:

– Can we use machine learning to extract breast tumor growth patterns that take into account patient variability and limited amount of data?
– Can we use machine learning to capture the peculiarities of tumor biology data by learning the underlying mathematical relations / functional dependencies of phenotypical transitions of cancer cells?
– Can we use machine learning to predict the volume of the breast affected by tumor (as for instance in DCIS) from limited and noisy timeseries of histopathologic data?

In the following sections we answer these motivating questions by demonstrating the capbilities of our model.

## 2 Materials and methods

In the current section we describe the underlying Machine Learning system along with relevant state-of-the-art models and datasets used in our experiments.

### 2.1 Models of tumor growth

Various tumor growth models have been proposed and are used to make predictions in cancer treatments planning [11]. In this work, we chose three of the most representative and typically used ordinary differential equations (ODE) growth models, namely Logistic, von Bertalanffy, and Gompertz, described in Table 2.1.

**Table 1.**
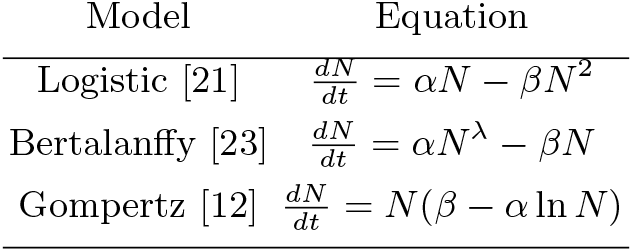
Tumor growth models in our study. Parameters: *N* – cell population size (or volume / mass thorough conversion [13]), *α* – growth rate, *β* – cell death rate, λ – nutrient limited proliferation rate, *k* – carrying capacity of cells.

### 2.2 Introducing our model

Our proposed solution is an unsupervised machine learning system based on Self-Organizing Maps (SOM) [14] and Hebbian Learning (HL) [3] used in combination in order to extract underlying mathematical relations among correlated timeseries describing tumor growth. In order to introduce our system, we provide a simple example in Figure 1. Here, we consider data from a cubic tumor growth law (3^*rd*^ powerlaw) describing the impact of sequential cytostatics dose density over a 150 weeks horizon in adjuvant chemotherapy of breast cancer [5]. The two input timeseries (i.e. the number of cancer cells and the irregular measurement index over the weeks) follow a cubic dependency, depicted in Figure 1a.

**Fig. 1.**
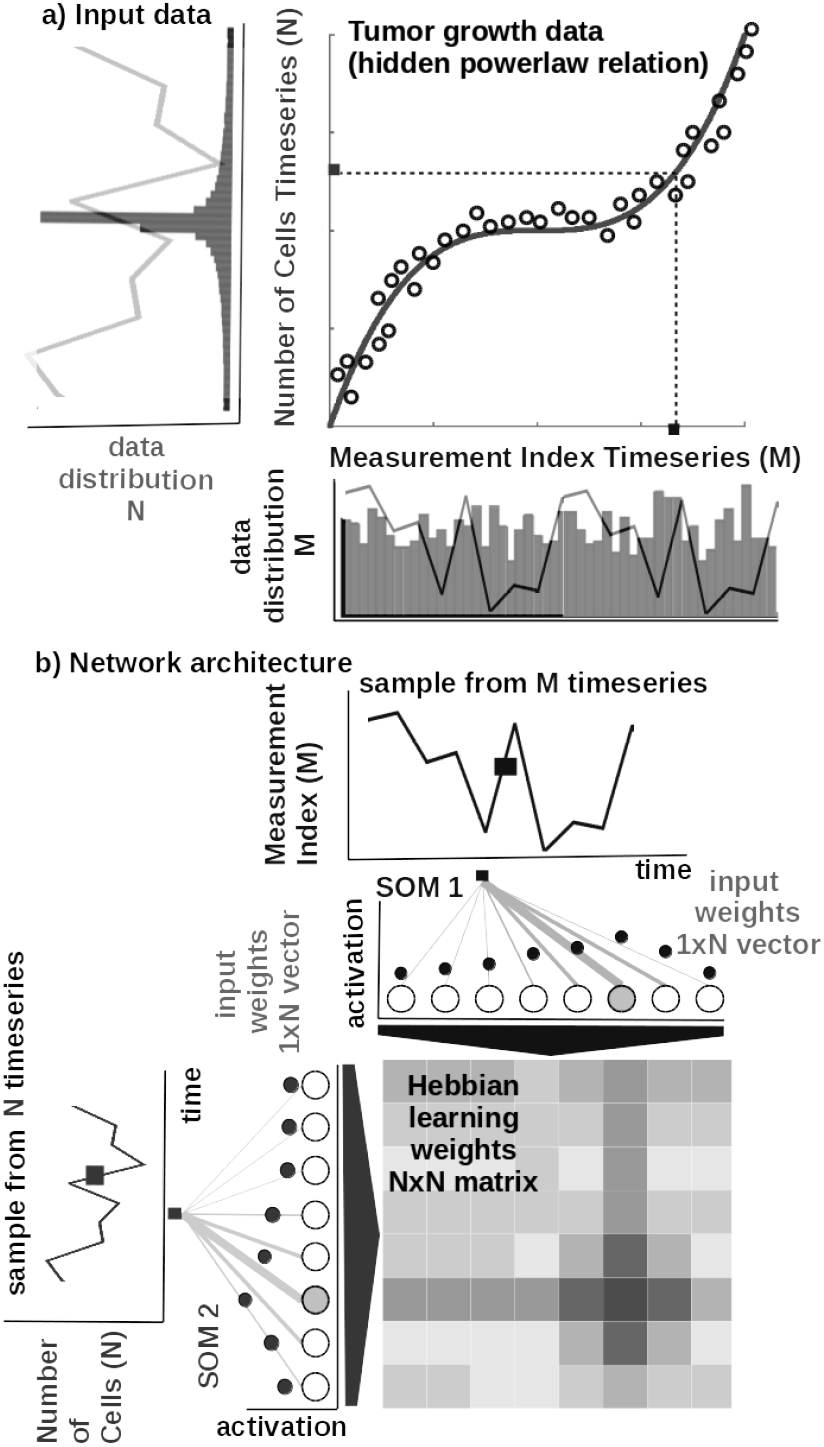
Basic functionality: a) Tumor growth data resembling a powerlaw (i.e. number of cells vs. measurement index). Data from [5]. b) Basic architecture of the system: 1D SOM networks with *N* neurons encoding the timeseries (i.e. number of cells vs. measurement index), and a *NxN* Hebbian connection matrix coupling the two 1D SOMs that will eventually encode the relation between the timeseries, i.e. growth curve.

#### Core model

The input SOMs (i.e. 1D lattice networks with *N* neurons) are responsible to extract the distribution of the timeseries data, depicted in Figure 1a, and encode timeseries samples in a distributed activity pattern, as shown in Figure 1b. This activity pattern is generated such that the closest preferred value of a neuron to the input sample will be strongly activated and will decay, proportional with distance, for neighbouring units. The SOM specialises to represent a certain (preferred) value in the timeseries and learns its sensitivity, by updating its tuning curves shape. Given an input sample *s^p^*(*k*) from one timeseries at time step *k*, the network computes for each *i*-th neuron in the *p*-th input SOM (with preferred value 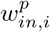 and tuning curve size 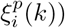 the elicited neural activation as

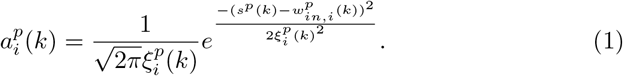

The winning neuron of the *p*-th population, *b^p^*(*k*), is the one which elicits the highest activation given the timeseries sample at time *k*

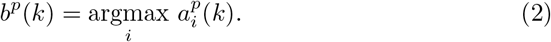

The competition for highest activation in the SOM is followed by cooperation in representing the input space. Hence, given the winning neuron, *b^p^*(*k*), the cooperation kernel,

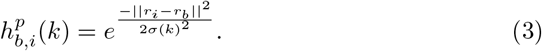

allows neighbouring neurons (i.e. found at position *r_i_* in the network) to precisely represent the input sample given their location in the neighbourhood *σ*(*k*) of the winning neuron. The neighbourhood width *σ*(*k*) decays in time, to avoid twisting effects in the SOM. The cooperation kernel in Equation 3, ensures that specific neurons in the network specialise on different areas in the input space, such that the input weights (i.e. preferred values) of the neurons are pulled closer to the input sample,

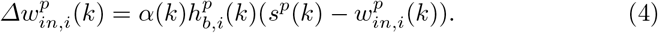

This corresponds to updating the tuning curves width 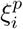 as modulated by the spatial location of the neuron in the network, the distance to the input sample, the cooperation kernel size, and a decaying learning rate *α*(*k*),

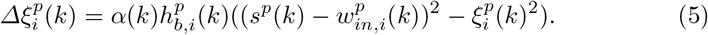

As an illustration of the process, let’s consider learned tuning curves shapes for 5 neurons in the input SOMs (i.e. neurons 1, 6, 13, 40, 45) encoding the breast cancer cubic tumor growth law, depicted in Figure 2. We observe that higher input probability distributions are represented by dense and sharp tuning curves (e.g. neuron 1, 6, 13 in SOM1), whereas lower or uniform probability distributions are represented by more sparse and wide tuning curves (e.g. neuron 40, 45 in SOM1). Neurons in the two SOMs are then linked by a fully (all-to-all) connected matrix of synaptic connections, where the weights in the matrix are computed using Hebbian learning. The connections between uncorrelated (or weakly correlated) neurons in each population (i.e. *w_cross_*) are suppressed (i.e. darker color) while correlated neurons connections are enhanced (i.e. brighter color), as depicted in Figure 2. Formally, the connection weight 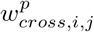 between neurons *i,j* in the different input SOMs are updated with a Hebbian learning rule as follows:

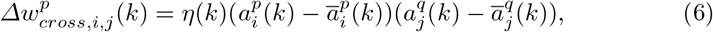

where

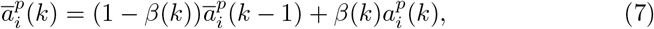

is a “momentum” like exponential decay and *η*(*k*), *β*(*k*) are monotonic (inverse time) decaying functions. Hebbian learning ensures that when neurons fire synchronously their connection strengths increase, whereas if their firing patterns are anti-correlated the weights decrease. The weight matrix encodes the co-activation patterns between the input layers (i.e. SOMs), as shown in Figure 1b, and, eventually, the learned growth law (i.e. relation) given the time-series, as shown in Figure 2.

**Fig. 2.**
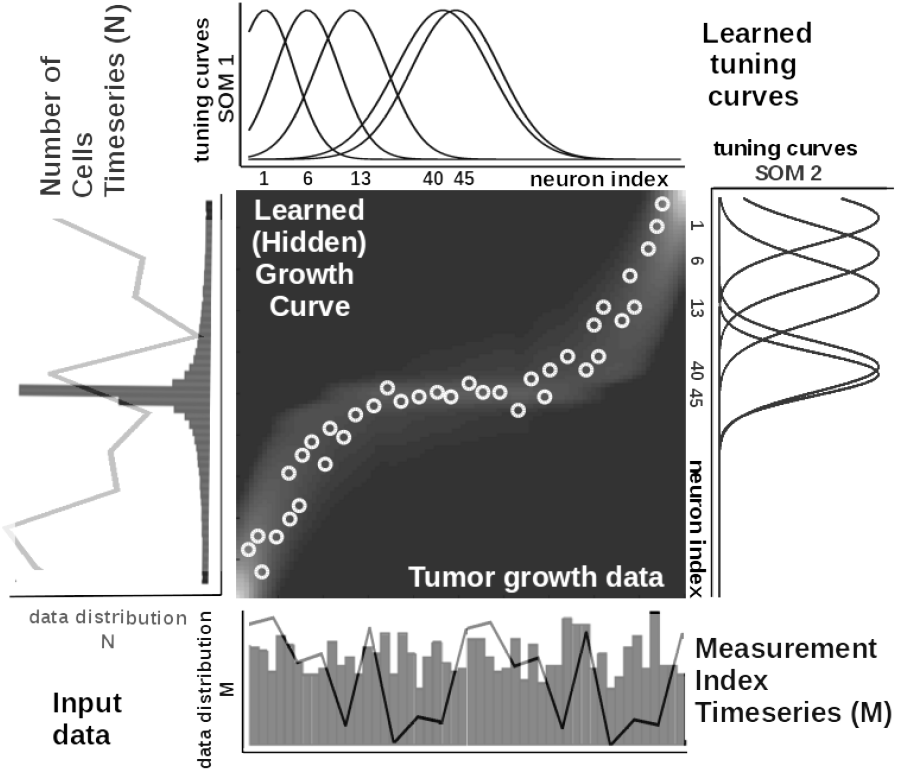
Extracted timeseries relation describing the growth law and data statistics for the data in Figure 1a depicting a cubic breast cancer tumor growth law among number of cells and irregular measurement over 150 weeks, data from [5]. Timeseries overlay on the data distribution and corresponding model encoding tuning curves shapes.

Self-organisation and Hebbian correlation learning processes evolve simultaneously, such that both the representation and the extracted relation are continuously refined, as new samples are presented. This can be observed in the encoding and decoding functions where the input activations are projected though *w_in_* (Equation 1) to the Hebbian matrix and then decoded through *w_cross_*.

#### Parametrization and read-out

In all of our experiments data from tumor growth timeseries is fed to the system which encodes it in the SOMs and learns the underlying relation in the Hebbian matrix. The SOMs are responsible of bringing the timeseries in the same latent representation space where they can interact (i.e. through their internal correlation). In all our experiments, each of the SOM has *N* = 100 neurons, the Hebbian connection matrix has size *NxN* and parametrization is done as: *α* = [0.01,0.1] decaying, *η* = 0.9, 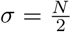 decaying following an inverse time law. We use as decoding mechanism an optimisation method that recovers the real-world value given the self-calculated bounds of the input timeseries. The bounds are obtained as minimum and maximum of a cost function of the distance between the current preferred value of the winning neuron and the input sample at the SOM level.

### 2.3 Datasets

For experiments we used publicly available clinical cancer datasets (see Table 2).

**Table 2.**
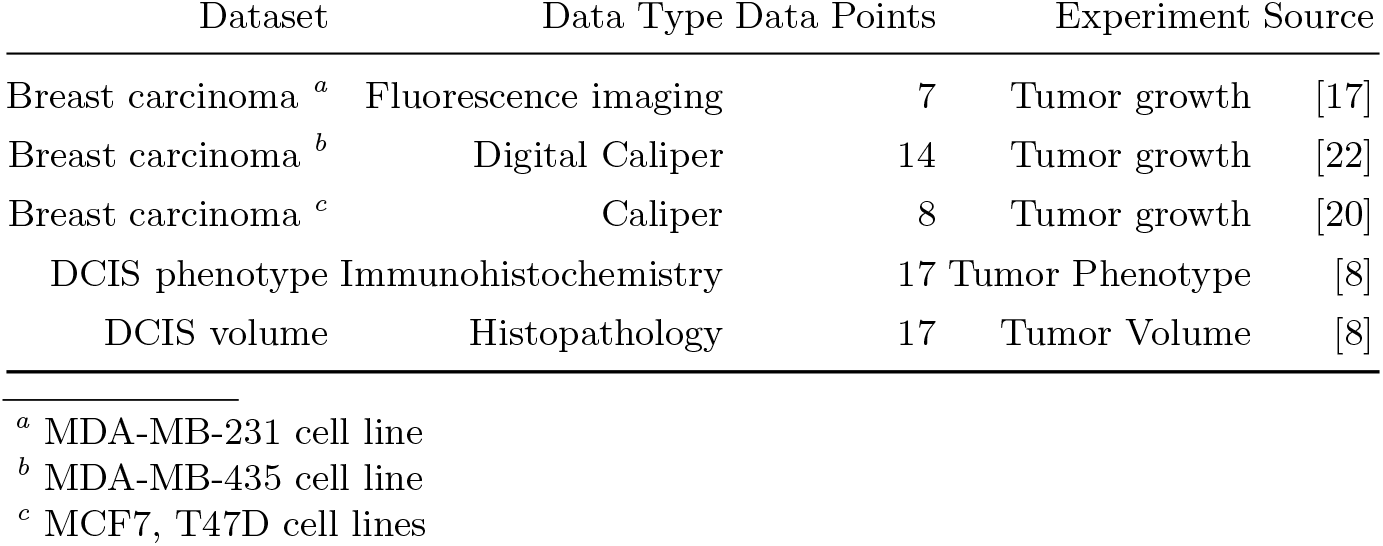
Description of the datasets used in the experiments.

### 2.4 Procedures

In order to reproduce the experiments and figures, the MATLAB ^®^ code and copies of all the datasets are available on GITLAB^4^.

For the tumor growth experiment: each of the three mechanistic tumor growth models (i.e. Logistic, Bertalanffy, Gompertz) and our model were presented the tumor growth data in each of the datasets (from [17,20,22]). When a dataset contained multiple trials, a random one was chosen.

For the tumor phenotype experiment our model was fed with timeseries of immunohistochemistry data and compared in accuracy against a mathematical model from [8].

For the tumor volume prediction experiment our model was fed with timeseries of histopathology data and compared in accuracy against a mathematical model and mammography data from [8].

#### Mechanistic models setup

Each of the state-of-the-art tumor growth models was implemented as ODE and integrated over the dataset length. We used a solver based on a modified Rosenbrock formula of order 2 that evaluates the Jacobian during each step of the integration. To provide initial values and the best parameters (i.e. *α, β*, λ, *k*) for each of the four models the Nelder-Mead simplex direct search was used, with a termination tolerance of 10*e*^−6^ and upper bounded to 500 iterations. Finally, fitting was performed by minimizing the sum of squared residuals (SSR).

#### Our model setup

For our model the data was normalized (interval [−1,1]) before training and de-normalized for the evaluation. The system was comprised of two input SOMs, each with *N* = 50 neurons, encoding the volume data and the irregular sampling time sequence, respectively. Both input density learning and correlation learning cycles were bound to 100 epochs.

## 3 Results

In the current section we introduce the results, discuss the findings, and demonstrate that our model can:

– extract breast tumor growth patterns that take into account patient variability and limited amount of data;
– learn the underlying mathematical relations / functional dependencies of phenotypical transitions of cancer cells;
– predict the volume of the breast affected by tumor (exemplified in DCIS) from limited and noisy timeseries of histopathologic data,

consistently with measurements in clinical setup.

### 3.1 Tumor growth curve extraction

In our first experiment, we explore the tumor growth curve prediction capability of our model across multiple breast cancer datasets. In all experiments, the state-of-the-art mechanistic models and our model were evaluated using Sum of Squared Errors (SSE), Root Mean Squared Error (RMSE), and Symmetric Mean Absolute Percentage Error (sMAPE). As our experiments demonstrate (Table 3), our model obtains superior accuracy in predicting the growth curve of the three types of tumors in three cell lines of breast cancer. Such a difference is supported by the fact that our model is a data-driven learning model that captures the peculiarities of the data and exploits its statistics to make predictions. The mechanistic models, on the other side, exploit biological knowledge and describe the dynamics of the physical processes, but fail in terms of versatility among tumor types.

**Table 3.** Evaluation of the tumor growth models.

| Evaluation Metrics |  |  |  |
| --- | --- | --- | --- |
| Dataset/Model | SSE | RMSE | sMAPE |
| *Breast<sup>a</sup> cancer [17]* |  |  |  |
| Logistic | 7009.6 | 37.4423 | 1.7088 |
| Bertalanffy | 8004.9 | 44.7350 | 1.7088 |
| Gompertz | 7971.8 | 39.9294 | 1.7088 |
| **Our model** | 119.3 | 4.1285 | 0.0767 |
| *Breast<sup>b</sup> cancer [22]* |  |  |  |
| Logistic | 0.2936 | 0.1713 | 0.1437 |
| Bertalanffy | 0.2315 | 0.1604 | 0.1437 |
| Gompertz | 0.3175 | 0.1782 | 0.1437 |
| **Our model** | 0.0977 | 0.0902 | 0.0763 |
| *Breast<sup>c</sup> cancer [20]* |  |  |  |
| Logistic | 3.0007 | 0.7071 | 1.0606 |
| Bertalanffy | 3.2942 | 0.8116 | 1.0606 |
| Gompertz | 3.1908 | 0.7292 | 1.0606 |
| **Our model** | 0.7668 | 0.3096 | 0.2615 |

<sup>a</sup> MDA-MB-231 cell line
<sup>b</sup> MDA-MB-435 cell line
<sup>c</sup> MCF7, T47D cell lines

### 3.2 Learning the phenotypical transitions of tumors

In order to demonstrate that our model can learn the mathematical relations describing the phenotypical transitions of tumors, we considered the study of 17 DCIS patients in [8]. In typical cancer phenotypic state space, quiescent cancer cells (Q) can become proliferative (P) or apoptotic (A). Non-necrotic cells become hypoxic when oxygen drops below a threshold value. Hypoxic cells can recover to their previous state or become necrotic [15]. Here we only focus on a 3-state sub-model (i.e. P, Q, A states). The transitions among theses sates are stochastic events generated by Poisson processes. Our model was fed with timeseries of raw immunohistochemistry and morphometric data for each of the 17 tumor cases (see [8], Tables S1 and S2) as following: cells cycle time *τ_P_*, cells apoptosis time *τ_A_*, proliferation index *PI* and apoptosis index *AI*. Using this input the system has to infer the mathematical relations for *α_P_*, the mean Q – P transition rate, and *α_A_*, the Q – A transition rate, respectively (see Figure 3). Their analytical form is:

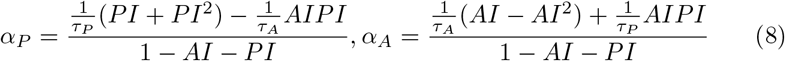

**Fig. 3.**
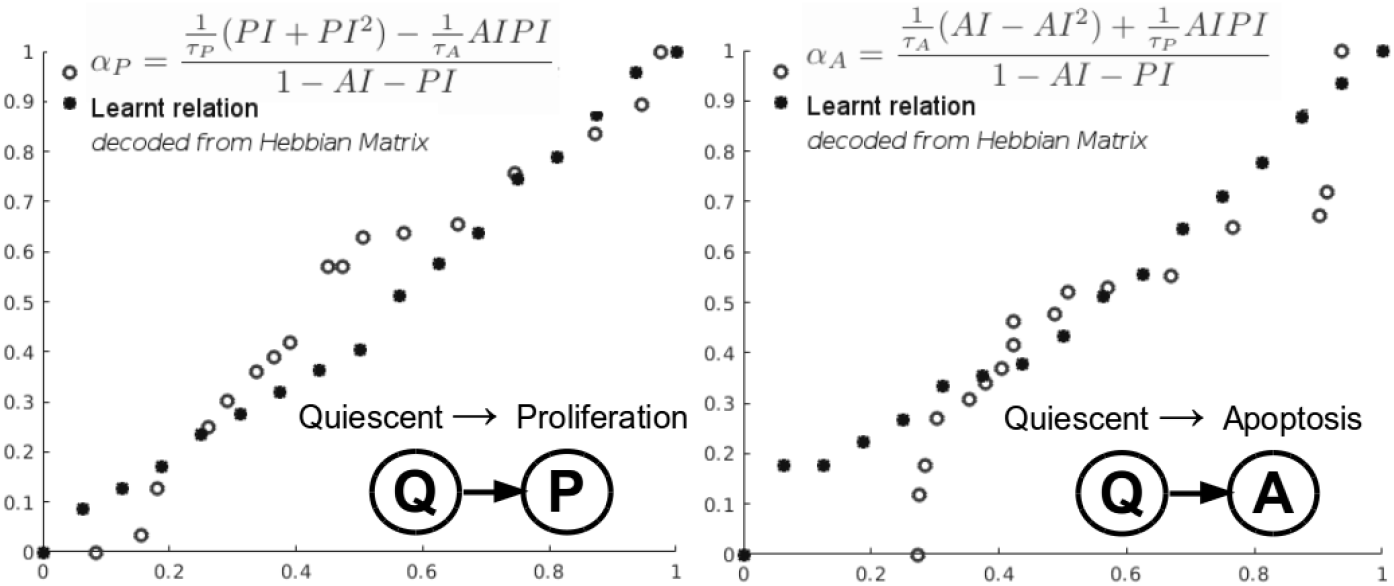
Learning cancer cells phenotypical states transitions mathematical relations.

Looking at the learnt mathematical relation describing the quiescent (Q) to apoptosis (A) and quiescent (Q) to proliferation (P) state transitions of cancer cells in Figure 3, we can see that our model is able to accurately recover the correct underlying mathematical function with respect to ground truth (clinically extracted and modelled Equation 8 from [15]): for Q to A transition *SSE* = 0.398, *sMAPE* = 0.131, *RMSE* = 0.153 and for Q to P transition *SSE* = 0.750, *RMSE* = 0.210, *sMAPE* = 0.172, respectively. Note that our system had no prior knowledge of the data distribution or biological assumptions, it simply learnt from the data the underlying relations using the neural processing described in section Materials and methods.

### 3.3 Prediction of tumor volume

In this section, we demonstrate that our model, without modification from the other experiments, can provably predict the surgical size of a tumor from pathology data on an individual patient basis.

Modelling surgical volume aims to elucidate the extent of the volume of tissue that must be surgically removed in order to (1) increase patient survival and (2) decrease the likelihood that a second or third surgery, or even (3) determine the sequencing with chemotherapy. We assessed this capability by demonstrating that our model can learn the dependency between histopathological and morphological data, such as nutrient diffusion penetration length within the breast tissue (*L*), ratio of cell apoptosis to proliferation rates (*A*) and radius of the breast tumor (*R*), in an unsupervised manner, from DCIS data of [8] (see Table 2). More precisely, the authors postulated that the value of *R* depends upon *A* and *L* following a “master equation”:

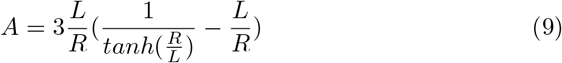

Its predictions, the study shown, are consistent with published findings that nearly 80% of in-situ tumors identified by mammographic screenings. Compared to ground truth, (Equation 9) our model was capable of extracting an accurate depiction of the growth pattern with *SSE* = 47.889, *RMSE* = 0.437 and *sMAPE* = 0.486, respectively. Remarkably, despite the lack of prior knowledge about the data and biological constraints, the model learned the growth curve and inferred the tumor evolution (see decoded relation in Figure 4). Our model is consistent with pathological/mammographic features predictions in [8] (Table 2). Our belief is that measuring such parameters at the time of initial biopsy, pathologists could use our model to precisely estimate the tumor size and thus advise the surgeon how much tissue (surgical volume) is to be removed.

**Fig. 4.**
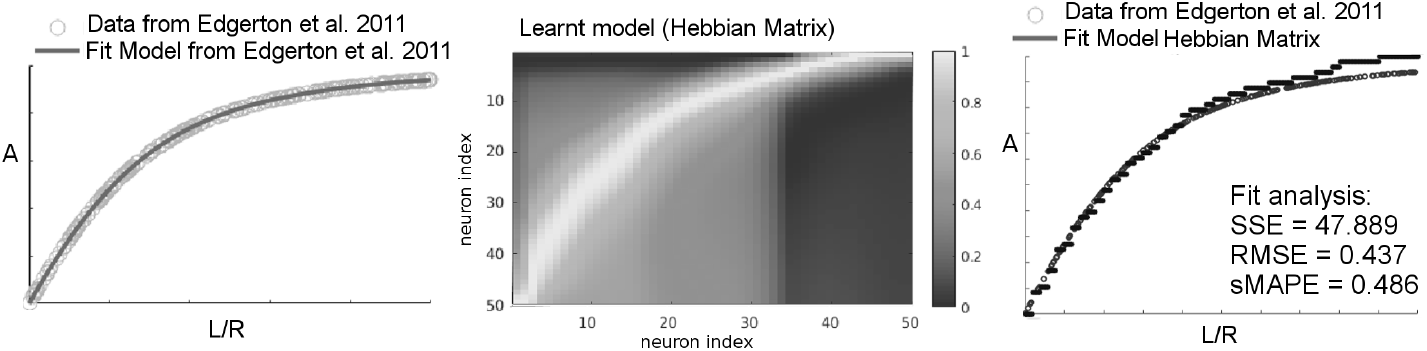
Upper panel: Modelled relation among *A* – the ratio of cell apoptosis to proliferation rates, *L* – the nutrient diffusion penetration length across tumor surgical volume, and *R* – the geometric mean tumor surgical radius, corresponding to Equation 9 on data from [8]. Middle panel: The learnt curve as visible in our model’s neural weight matrix (i.e. Hebbian connection matrix). Lower panel: Evaluation of the learnt relation against the analytical (ground truth) model in Equation 9. *In the Hebbian matrix brighter tones code higher values, darker tones lower values*.

## 4 Conclusion

There is a significant unfulfilled need for more accurate methods to assess the volume of a clinically diagnosed breast cancer before planning surgery or therapy. Tackling this need, we proposed a versatile unsupervised learning system that is able to extract breast tumor growth patterns that take into account patient variability and limited amount of data from 3 different breast cancer cell lines and overcome 3 state-of-the-art models. In a second experiment, using the same computational substrate, our model learned the underlying mathematical relations describing the phenotypical transitions of the cancer cells, consistent with clinical data. Finally, our model predicted the volume of the breast affected by tumor from limited and noisy timeseries of histopathologic data from 17 cases of DCIS. This suite of experiments prove the versatility of the model and propose it as a candidate for more accurate assessment of surgical volume, which could improve the success of complete excision of breast tumors.

## Footnotes

4 https://gitlab.com/akii-microlab/icann-2020-bio

## Notes

### Competing Interest Statement

The authors have declared no competing interest.

